# Ridge-assisted Micro Positioning of Cells and Particles in a Microchannel

**DOI:** 10.1101/2025.09.26.678893

**Authors:** Adriana Payan-Medina, Victor R. Putaturo, Sajad R. Bazaz, Quinn E. Cunneely, Varun Kulkarni, Mehmet Toner, Avanish Mishra

## Abstract

The capability to deterministically and precisely control the lateral position of focused cell streams in a microchannel provides diverse opportunities for biomedical applications, including cell concentration, imaging flow cytometry, and cell-based liquid biopsy platforms. Inertial microfluidics introduced various high-throughput strategies for ordering particles and cells within microchannels. However, existing approaches do not permit precise control over the lateral focusing position of cells. By leveraging engineered microvortices generated by channel ridges and fluidic lift forces, we present a microfluidic method for deterministically positioning focused cell streams to desired lateral streamlines. Unlike inertial focusing, the ridge-assisted micro positioning (RAMP) is independent of particle size and flow rate, enabling polydisperse particles (10 to 30 μm) to be focused to the same streamline across a broad range of flow rates (250–1000 μL/min). The focusing position can be precisely adjusted by altering ridge placement and geometry, achieving both small (5 μm) and large (≥15 μm) lateral shifts in a controlled manner. Applying the RAMP concept, we developed clog-free microfluidic cell concentrators that can enrich single cells and clusters of cells to desired concentration factors and demonstrate the ability to focus particles in undiluted whole blood. Together, these results establish RAMP as a versatile platform for precise cell positioning in complex biological fluids.

**Teaser:** Use of designer vortices produced by ridges allows cells to be focused in user-defined streamlines across a microchannel over a wide range of flow rates.

## Introduction

Microfluidic devices are invaluable tools for analysing complex biofluids such as blood, pleural, or peritoneal fluids for disease monitoring and diagnosis (*4–6)*. These samples vary substantially in composition across patients and contain heterogeneous cell populations (*7*). For such analyses, precise positioning of cells within a microchannel is critical. Flow cytometry, cell sorting, and sensing all rely on consistent placement of cells at streamlines where sorting forces or sensing fields are optimal, thereby improving measurement sensitivity and accuracy (*8*).

Conventional flow cytometry systems achieve this two-dimensional alignment using coflowing buffer sheath streams to confine cells at the centre of the optical path (*9*). Sheath streams used for alignment in imaging flow cytometry are designed for cells to be tightly confined in all three dimensions to reduce dropout events (*10, 11*).

Inertial microfluidics has provided an elegant strategy for high-throughput cell ordering with regulated spacing (*12–17*) (**Table S1**)12/22/2025 5:00:00 PM. However, the relative focusing position is determined by the channel geometry and cannot be independently controlled. Thus, a key unmet need has been the development of a microfluidic system that achieves deterministic focusing and permits precise (micron-level) and adjustable control of lateral cell positioning. In this study, we introduce a ridge-assisted micro positioning (RAMP) platform that enables polydisperse particles to be focused into single streamlines. Critically, the lateral position of the focused stream can be predictably adjusted by modifying the ridge placement and geometry, enabling micron-level control from the channel sidewall to its centreline (**Figure 1**). This tunability distinguishes RAMP from previous inertial focusing approaches and establishes a versatile foundation for a range of biomedical applications.

**Fig. 1.**
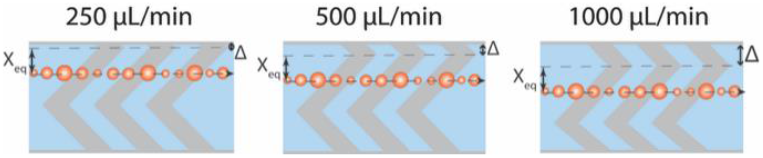
Ridge-assisted micro positioning (RAMP) achieves controllable focusing of polydisperse particle streams. Channels with RAMP can precisely adjust polydisperse particles’ focusing position as the ridge structure location is shifted (ridge shift indicated with *Δ*, while particle focusing position, X_eq_, remains constant) at several high-throughput flow rates. Thus, shifts in central ridge position, *Δ*, result in proportional, micron-level shifts in particle focus position.

## Results

### RAMP Design and Focusing Principles

The focusing mechanism for RAMP can be attributed to secondary flows generated by the ridges incorporated into the microchannel (**Figure 2A-D**). In a rectangular channel without microstructures, particles migrate to equilibrium positions near the long face of the channel, where shear gradient and wall-induced lift forces oppose one another (**Figure 2E**) (*1, 18*). The introduction of ridges disrupts this balance by inducing anisotropic pressure gradients along their inclined surfaces, which drive fluid recirculation parallel to the ridge direction (*19*) (**Figure 2B**). This recirculating flow forms a transverse vortex, exerting drag forces on suspended particles. In combination with inertial lift forces, these drag forces guide particles toward stable focusing positions (**Figure 2C-D**) (*15*).

**Fig. 2.**
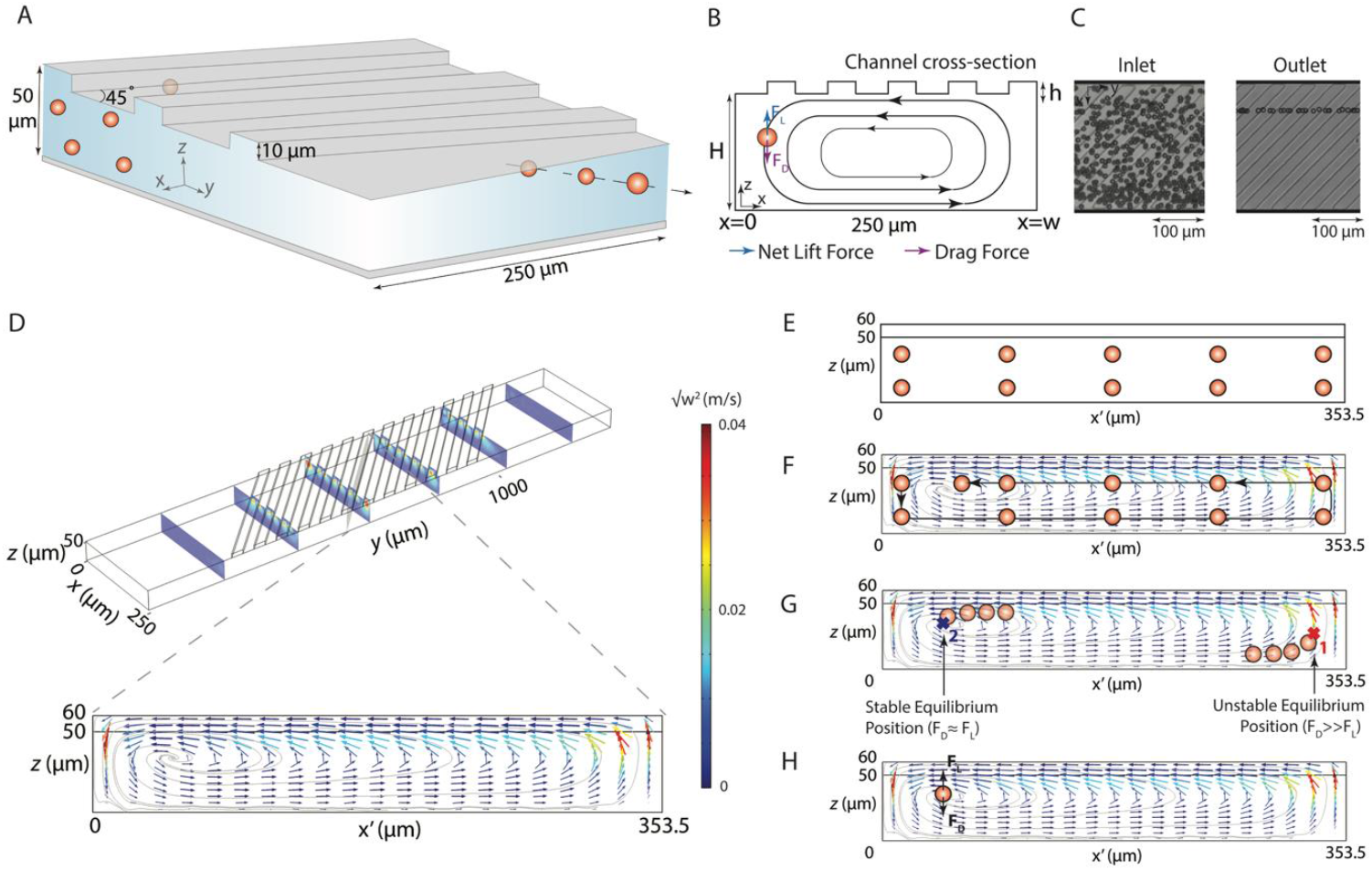
RAMP device schematics and fluid flow simulation show relevant geometric features and focusing mechanism. (**A**) A schematic representation of a RAMP device showing the focusing effect observed in a microchannel due to a transverse vortex caused by ridges. (**B**) The microvortex exerts a drag force on particles. When drag and lift forces are balanced, particles assume a stable equilibrium position. (**C**) High-speed microscopy images show dispersed 10 μm particles at the inlet and focused 10 μm particles at the outlet at 500 μL/min (Re = 50.4). (**D**) Finite element modelling results illustrate transverse velocity vectors and flow recirculation regions with color-mapped velocity magnitudes. (**E-H**) Mechanism of particle focusing. In a rectangular channel without ridges, particles undergo inertial ordering along the long faces of the channel (**E**). Transverse vortex, produced by ridges, sweeps particles (**F**). Theoretically, two equilibrium positions are possible (**G**), one near the center of the vortex (1) and another near a side wall, away from the center of vortex (2), but drag force *F*_*D*_ is 10-fold stronger near position 2, causing particles to be swept away from position-2 to a stable equilibrium position-1, near the stagnation region at the center of the vortex where *F*_*D*_ is feeble (**H**).

The approximate lift and drag force equations for particle trapping are described in the **Supplementary Text** with **Equations S1** to **S4**. A simplified ratio of these two forces (*F*_*L*_*/F*_*D*_) (**Equation S4**) demonstrates that particle focusing is dictated by the confinement ratio, *a/H*, where *a* is the particle diameter and *H* is the channel height, and a geometrical nondimensional factor, *α* = *h/*2*H*, where *h* is the ridge height. The strength of the transverse flow can be further tuned with the ridge geometry (angle (*θ*), height (*h*), and wavelength (*λ*)), channel dimensions (Width (*W*) and height (*H)*), and Reynolds number (*Re*) (*19*). Based on this analysis, the ridge and channel dimensions were adjusted to have a low *α* and high confinement ratio (**Figure 2**), resulting in a balance of lift-to-drag forces (*F*_*L*_ ≈ *F*_*D*_). The resulting focusing effect shows 10 μm particles focused near the channel sidewall at the outlet, contrasting with the dispersion observed at the inlet (**Figure 2C**). In this case, microchannels are 250 μm wide (*W, x*-direction), 50 μm deep (*H, z*-direction), and *Re* is 50.4 while the particle confinement ratio is 0.2. The height of ridges is 10 μm, with a 60 μm wavelength. Ridges are angled at 45º with respect to the long axis of the channel (**Figure 2A**).

Computational fluid dynamics (CFD) simulations confirmed the ridge-induced vortex that sweeps the particles to a stable focusing position (**Figure 2F** and **Figure S1**). These computational results show the microvortex in a cross-section parallel to the ridge, as illustrated in **Figure S2**. In a low-aspect-ratio rectangular channel with ridge-induced vortices, at moderate to high *Re*, theoretically, two focusing positions, 1 and 2, are possible (**Figure 2G**), each near the side channel wall. However, near the distant side wall (position-1), the drag force, *F*_*D*_, is 10-fold stronger than the lift force, *F*_*L*_, (*F*_*L*_*/F*_*D*_ ≅ 0.1027) making it an unstable equilibrium position. Here, *F*_*L*_*/F*_*D*_ was calculated using **Equation S2** for a flow rate of 500 µL/min (Re = 50.4). This is an approximate analysis based on an empirical relationship for lift force and velocity field estimated using CFD modeling, but it provides important insights into focusing behavior. Thus, particles at the unstable equilibrium position are swept away from this position to a stable equilibrium position, near the center of the vortex (position-2), where the *transverse* components of the velocity (*u* and *w*) are negligible, allowing balancing of the lift and drag forces (*F*_*L*_ *= F*_*D*_) (**Figure 2H**). This is also supported by the fact that, based on CFD simulations, the stagnation region at the center of the vortex is expected at *z* = 34 µm, while experimentally, it is observed at *z* = 31.5 ± 0.6 µm. Thus, the stagnation region at the center of the vortex can indicate the stable trapping region.

### RAMP facilitates polydisperse particle focusing across a wide flow rate range

We next examined whether RAMP could maintain stable particle focusing across a broad flow-rate regime. Results demonstrated that particles ranging from 10 to 30 μm in diameter could be consistently focused at flow rates between 250 and 1000 μL/min (**Figure 3**). **Figure 3A** shows streak images of 10-micron particles at flow rates ranging from 250 to 1000 μL/min (*Re* = 25.2 to 100.8). In a 250 µm wide cross-section, transverse to flow, 10 μm particles focused between 52.2±0.5 to 57.8±0.7 μm (**Table S2**). A frequency distribution plot of particle focusing position illustrates that the equilibrium position remains consistent across the flow rate range with negligible dispersion (**Figure 3B**). CFD simulations for each flow rate tested also demonstrate the formation of flow-rate-independent transverse recirculation regions (**Figure S3**), and the stagnation region does not change its position. A high-speed camera video shows 10 μm particles precisely aligned in a streamline at 500 μL/min (**Movie S1**). These computational and experimental results demonstrate that RAMP can focus particles in a flow-rate-insensitive manner in the *x*-*y* plane, with minimal dispersion in the equilibrium position of particles.

**Fig. 3.**
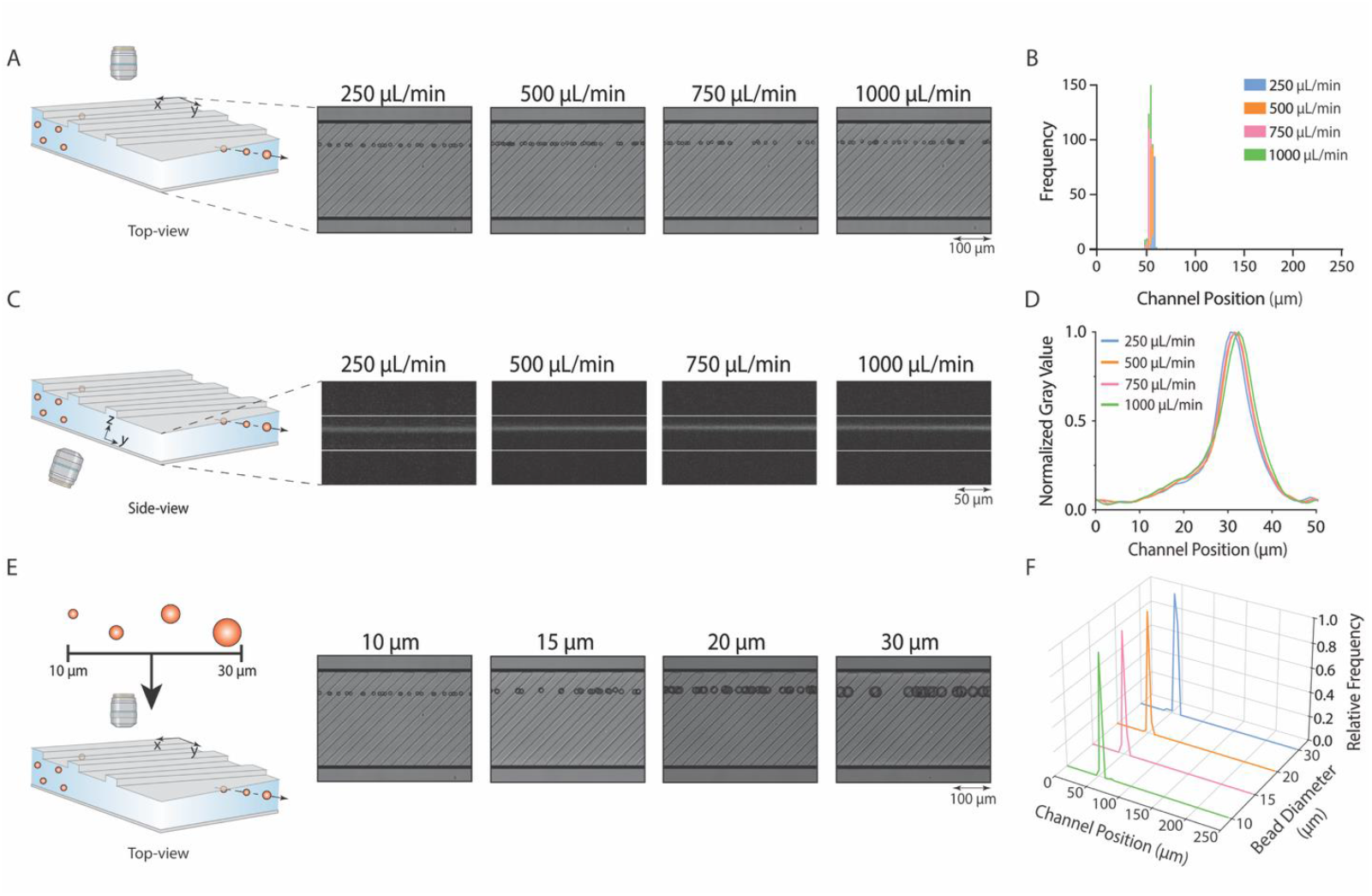
Microparticle positioning device with straight ridges enables focusing of particles across a wide range of flow rates in *x, y*, and *z* directions with particles of different sizes. **(A**) High-speed streak imaging shows 10 μm particles maintaining their focus position when flowing at several flow rates. (**B**) A frequency distribution diagram of particle centroid positions quantifies dispersion in focusing position at flow rates of 250 μL/min (n = 96, *Re* = 25.2), 500 μL/min (n = 166, *Re* = 50.4), 750 μL/min (n = 111, *Re* = 75.6), and 1000 μL/min (n = 114, *Re* = 100.8). (**C**) Fluorescent microscopy images show side-view images of 10 μm particles in the *y-z* plane. (**D**) Fluorescent intensity profile of the focused streamline shows particles focusing in the same *z*-plane for flow rates of 250 to 1000 μL/min (n = 20 profiles per flow rate). (**E**) High-speed streak imaging shows a top-view of particles with diameters ranging from 10 to 30 μm aligned at the same position in the channel. (**F**) Frequency distributions of particle centroid positions for 10 μm (n = 104), 15 μm (n = 100), 20 μm (n = 69), and 30 μm (n = 102) particles, showing focus position data for a flow rate of 250 μL/min. Data for other flow rates follow a similar trend (**Figure S4**).

Next, we examined the effect of varying flow rate on particle focus position in the *z*-direction (**Figure 3C**). In the *z*-direction, the particle focusing positions for each flow rate tested ranged from 30.6±0.6 to 32.4±0.5 μm, as measured from the bottom of the channel (**Figure 3D, Table S3**). These results demonstrate that RAMP achieves single-streamline focusing in all three dimensions (*x, y*, and *z*), a capability not afforded by conventional inertial focusing devices, such as rectangular channels (*20*), spiral focusing (*21*), or asymmetric serpentine channels (*22*), which typically yield two focusing positions in the *z*-direction. This 3D single-line focusing feature is particularly useful for imaging flow cytometers, as it minimizes out-of-focus events and enables consistent, reliable quantification of cells.

To ensure that RAMP’s precise flow-rate insensitive focusing capabilities extend to polydisperse particles, we tested 10 μm, 15 μm, 20 μm, and 30 μm particles at flow rates ranging from 250 to 1000 μL/min (*Re* = 25.2 to 100.8 and confinement ratios of 0.2, 0.3, 0.4, and 0.6). High-speed streak images of particles at 250 μL/min show particles focusing in approximately the same position (**Figure 3E-F**). Particles of various diameters (10 μm, 15 μm, 20 μm, and 30 μm) had average focus positions ranging from 54.2±1.3 to 59.5±2.3 μm (**Table S2**). A high-speed camera video shows 10 and 30 μm particles flowing at 500 μL/min through a RAMP device (**Movie S2**), creating an identical streakline. This consistency in polydisperse particle focusing was also observed at flow rates of 500, 750, and 1000 μL/min (**Figure S4, Table S2**). These analyses demonstrate that a RAMP device can tightly focus polydisperse particles in a wide flow rate regime with a minimal variation in the focus position.

### RAMP for precise adjustment of particle focus position

Beyond focusing polydisperse particles over a wide flow regime, RAMP provides precise control of particle positioning across the channel cross-section. Shifting the position of the ridge structure within the channel causes proportional shifts in particle focus position (**Figure 4A**, regions 4 and 5). To demonstrate this capability, we fabricated two sets of RAMP devices designed for either small (∼5 μm) or large (15 μm and 30 μm) focus position shifts. In these devices, microparticles are first aligned on both sides of a chevron microstructure within the channel, arranging particles into two streamlines (**Figure 4A**, region 2). Following the chevron region, a region with a central ridge structure joins these streamlines into one near the channel center (**Figure 4A**, region 3). Once particles are concentrated in a common region within the channel, small or large shifts in focus position are created by shifting the central ridge (**Figure 4A**, region 4-5, and **Figure S5**).

**Fig. 4.**
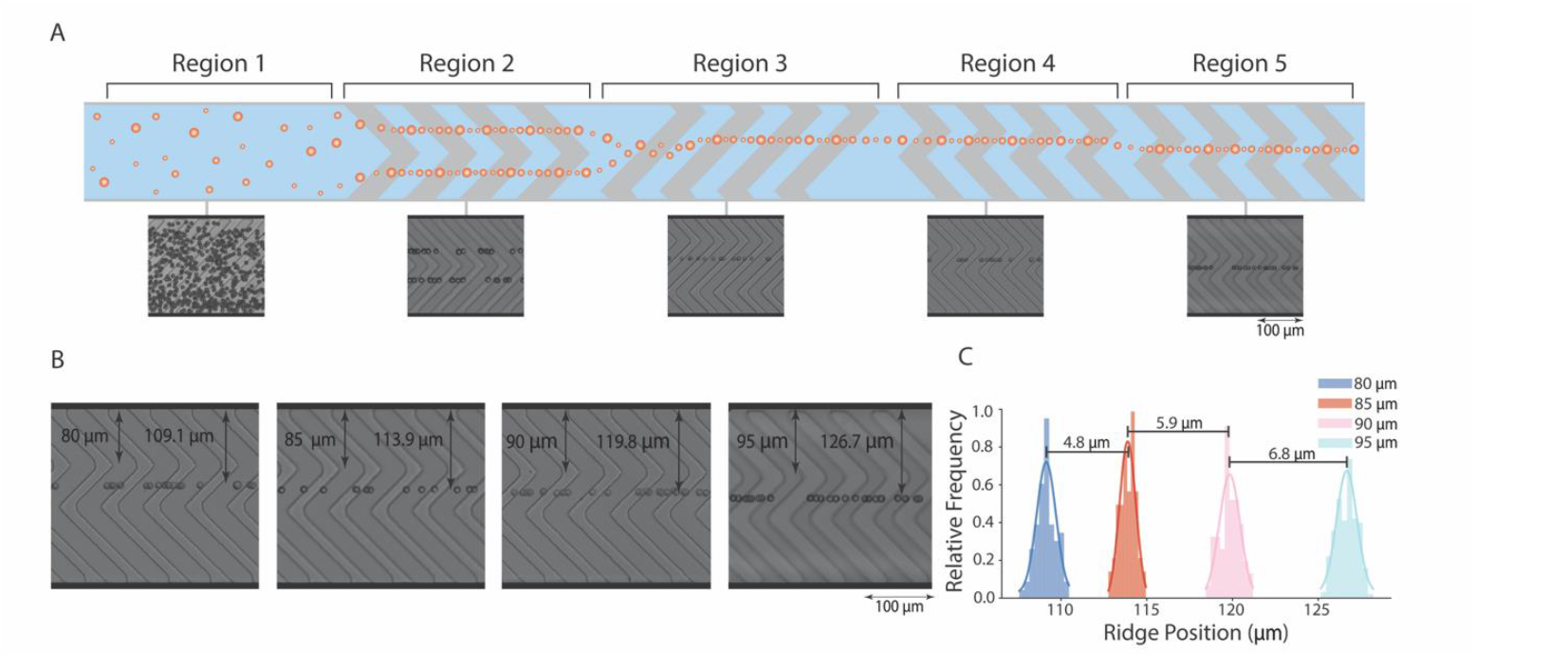
Ridge-assisted micro positioning device can achieve shifts in particle focus position through proportional shifts in ridge position. (**A**) Particles at the inlet (region 1) are aligned with symmetrical ridges in region 2 that bring particles into two streamlines. Region 3 uses a long, single ridge to align particles in a single line near the center of the channel. In region 4, particles flow into a region with an alternative central ridge structure. Between regions 4 and 5, the central ridge’s location is shifted to create proportional shifts in particle focus position. (**B**) High-speed streak images with reference distances show that the focused particle stream shifts proportionately as a central ridge is shifted 80 to 95 μm from the channel side wall. (**C**) A frequency distribution of particle centroid positions at ridge shifts of 80 μm (n = 83), 85 μm (n = 65), 90 μm (n = 58), and 95 μm (n = 314).

Shifts in the location of the central ridge create a proportional shift in the particle focus position. As the central ridge structure is shifted in steps of 5 μm from 80 μm to 95 μm in the mylar mask design (**Figure 4B**), secondary flow created by this structure caused focused particle streams to shift in steps of 4.8±0.7 μm, 5.9±0.8 μm, and 6.8±0.9 μm, respectively (**Figure 4B** and **Table S4)**. Slight variations from the targeted 5 μm shift in design to experimentally achieved shifts of 4.8 μm to 6.8 μm can be attributed to SU-8 photolithography tolerances, where features from the digital design are not exactly reproduced in SU-8, especially in this case, as a mylar mask was used, which has a slightly lower resolution than the glass chrome masks. A high-speed imaging video of 10 μm particles flowing through devices with center ridge structures at 80 μm and 90 μm further illustrates this controlled shift (**Movie S3**). Similarly, as a central ridge structure is shifted in larger steps of 15 μm and 30 μm, proportionate focus position shifts of 17.1±0.9 μm and 29.6±1.2 μm are achieved (**Figure S5** and **Table S4**). To demonstrate that this concept also works with polydisperse particles, we tested a mixed population of 10 μm and 15 μm particles flowing through the RAMP device (**Figure S6**). As expected, 5 μm shifts in ridge location from 80 μm to 90 μm resulted in particle focus position shift steps of 5.7±1.3 μm and 6.4±1.3 μm, respectively (**Figure S6B** and **Table S4**), showing the remarkable lateral positioning capability of this approach for particles of various sizes. Again, slight deviations from targeted design steps of 5 μm can be attributed to tolerances in SU-8 microfabrication. These results confirm that RAMP enables deterministic and finely tunable lateral positioning for monodisperse and polydisperse particle populations.

Another versatile feature of the RAMP platform is that the focus position is weakly dependent on channel cross-sectional width for low aspect ratio channels, *W/H>>*1 (**Figure 5**). Particles flowing through channels with an aspect ratio of 2.5 to 1.4 focus in the same equilibrium position (labeled as X_eq_) (**Figure 5A-B**). However, larger aspect ratio channels (*W/H* = 1.1) create equilibrium positions closer to the channel center (**Figure 5A-B**). Finite element simulations show the transverse velocity field in a cross-section parallel to ridge microstructures for three channels, with each successive channel half the width and half the flow rate of the one above it, starting from 250 µm and 500 μL/min. Channel and ridge heights remain constant at 50 μm and 10 μm, respectively (**Figure 5C**). These simulations correspond to channel widths of 250, 125, and 62.5 µm, from top to bottom. For *W/H*>>*1*, the flow field and stagnation point remain the same, contributing to the equilibrium position consistency observed experimentally; however, when *W*≈*H*, the stagnation point shifts toward the channel center. This analysis shows that the equilibrium position remains stable in low aspect ratio channels and provides an understanding of channel geometry design considerations.

**Fig. 5.**
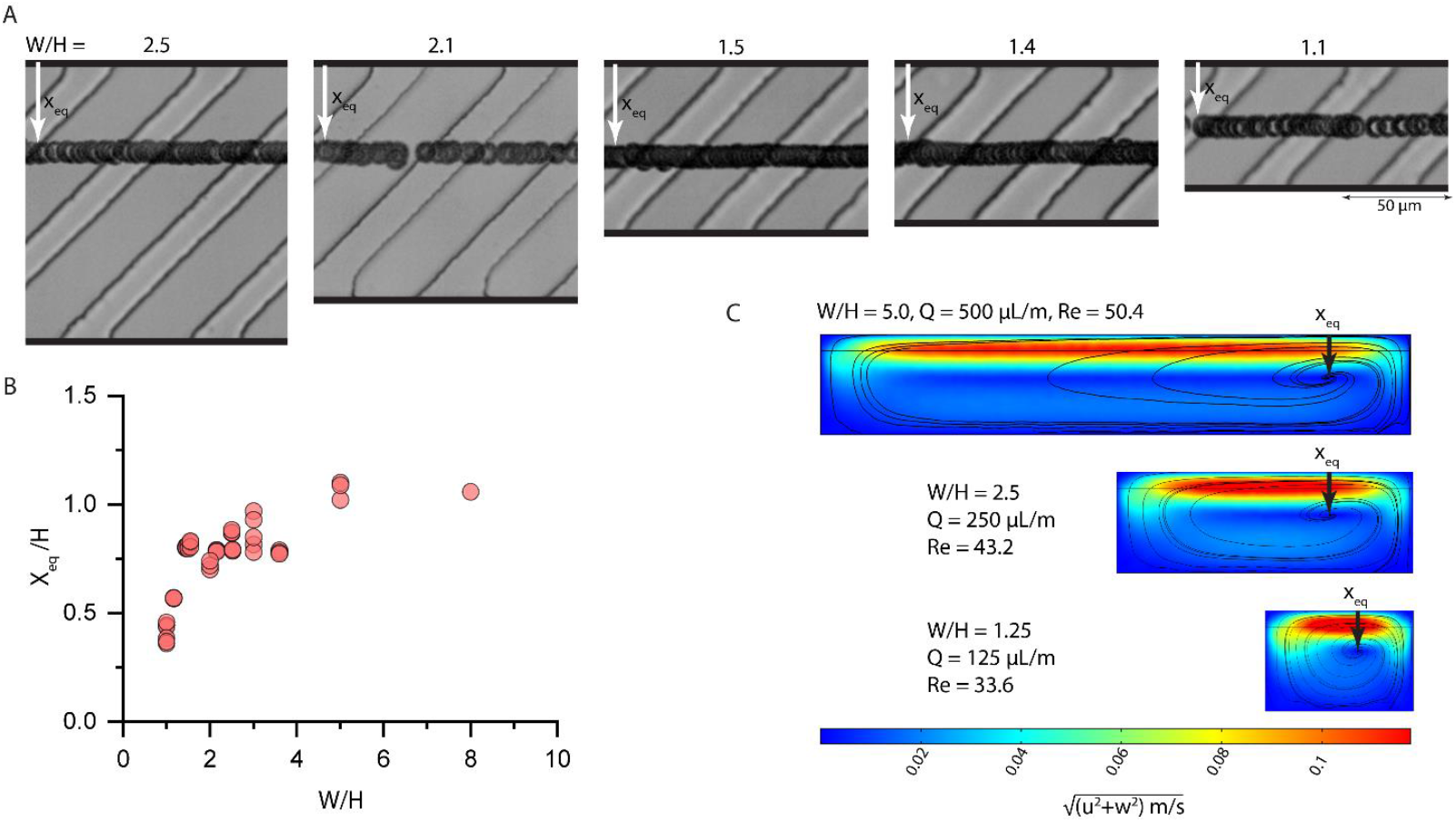
The RAMP focusing mechanism is independent of channel width for low aspect ratio channels (*W/H*>>1). (**A**) The equilibrium focus position (X_eq_) remains constant as 10 μm particles flow through channels of varying widths (unless *W* ≈ *H*). (**B**) A scatter plot displays average focus positions of particles flowing in channels of various widths. (**C**) Finite element fluid flow simulations show color-mapped transverse velocity magnitudes and black curves representing streamlines for three channels of varying widths and flow rate. In these simulations, the position of the stagnation region where trapping occurs remains unchanged (labeled with an arrow as X_eq_), unless *W* ≈ *H*.

### RAMP for cell concentration

RAMP’s practical benefits for sample preparation are exemplified through its use in cell concentration. Existing microfluidic concentrators often utilize free-standing posts that trap cellular debris or neutrophil extracellular traps (NETs). As a result, they are susceptible to clogging and compromised device performance (*21*). Using the RAMP concept, we developed a concentrator utilizing various ridge configurations to achieve efficient and reliable concentration of polydisperse particles without using any free-standing posts. In this concentrator device, we exploit three key features of RAMP: 1) focusing position consistency despite changes in the Reynolds number (25 to 100) or average velocity of the fluid flow as the cross section increases, 2) the alignment of polydisperse cells to identical focusing positions, and 3) the ability to align particles into specific streamlines by changing the ridge location. A suspension of cells or particles enters the concentrator (**Figure 6A**, region 1), where cells are progressively organized through three alignment regions. First, a 100 μm wide and 50 μm deep channel with 60 µm in wavelength chevron microstructures aligns cells into two symmetrical streamlines (**Figure 6A**, region 2). Then, the two symmetrical streamlines converge into a single streamline as cells enter a third region with single ridge microstructures (**Figure 6A**, region 3). Once aligned, cells retain their focus position, even as the bottom region of the channel is expanded with the siphoning angle (*θ*), allowing cell-free fluid to be efficiently removed in the siphoning layer (**Figure 6A**, region 4). High-speed images of particles in regions 1-4 of the concentrator demonstrate the stepwise concentration principle developed using RAMP (**Figure S7A**). We can achieve desired concentration factors by increasing the siphoning angle. We tested concentrators with a mixture of white blood cell and circulating tumor cell (MGH-BRx-142) population (**Figure 7B**) and a mixture of 10 and 20 μm particles (**Figure S7B**). When mixed cell populations were processed through the concentrators, concentration factors of 3.1X, 5.4X, 7.7X, 10.0X, and 12.3X yielded average cell recovery rates of 97.6±1.1%, 99.5±0.2%, 98.4±0.2%, 95.1±0.5%, and 96.8±0.8%, respectively (**Figure 6C, Table S5**). For polydisperse particle suspensions, recoveries remained consistently high at 99.7±0.2%, 99.8±0.1%, 99.7±0.2%, 99.0±0.1%, and 99.6±0.1% across the same concentration factors (**Figure S7C, Table S5**). Overall, particles and cells exhibited excellent yields, resulting in more than 95% cell recovery for all tested concentrator conditions, including polydisperse suspensions and cell populations. Minor cell losses in the mixed cell population can be attributed to dead cells or debris in the suspension, labeled in **Figure 6B**. High-speed video imaging further confirmed on-chip concentration of bead and cell suspension in RAMP concentrators at concentration factors of 3.1X and 12.3X (**Movies S4** to **S7**).

**Fig. 6.**
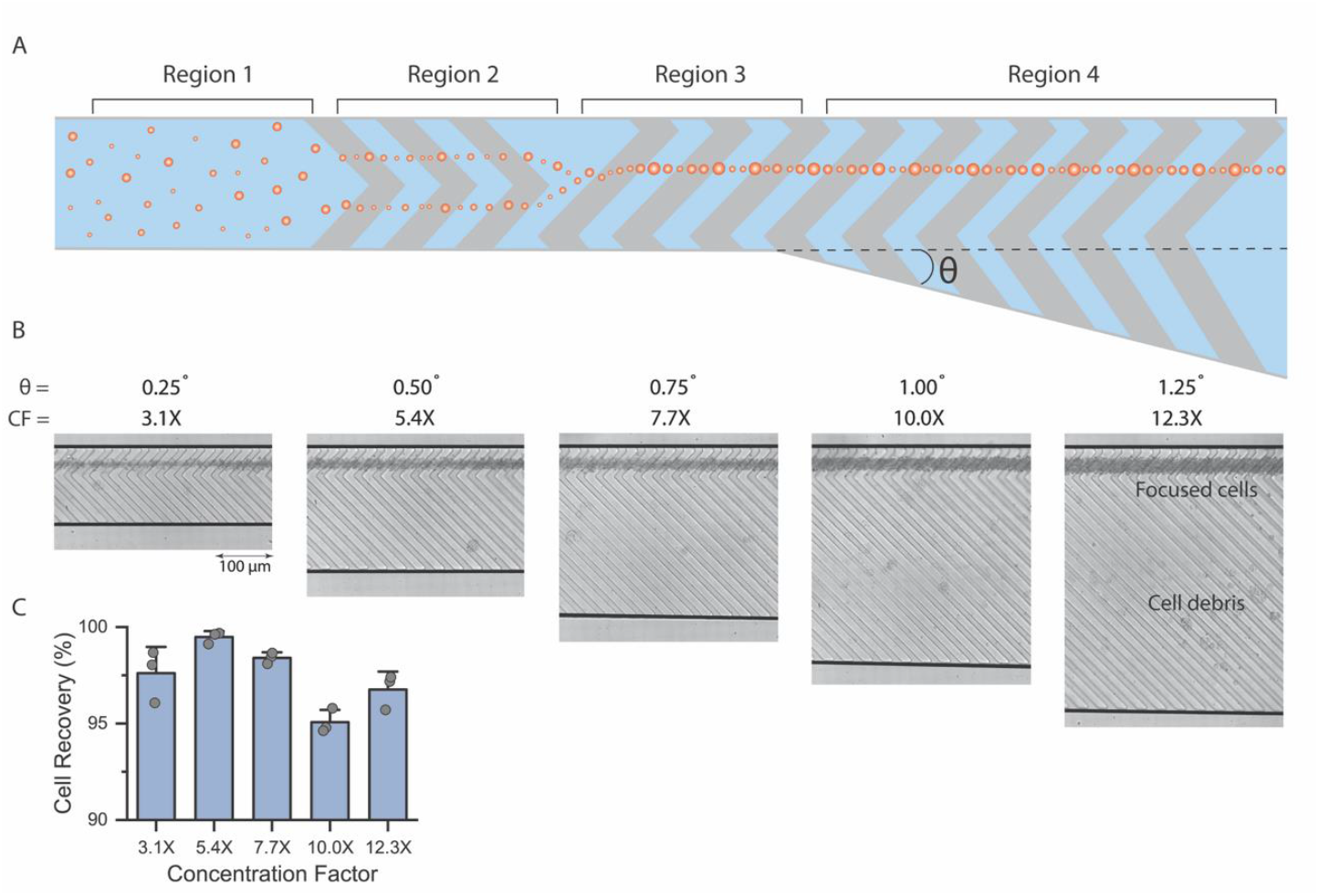
Microfluidic concentration of cells and polydisperse particles using RAMP. (**A**) A schematic representation of a concentrator with RAMP illustrates the focusing mechanism applied for concentration: region 1 shows particles entering the device, region 2 shows focusing of particles with symmetrical ridges that bring particles to the center of the channel in two streamlines, and region 3 is a long, single ridge that aligns particles in a single line near the center of the channel. Region 4 features a siphoning region, which removes cell-free fluid, concentrating cells. (**B**) RAMP concentrators enrich a mixture of white blood cells and circulating tumor cells at concentration factors of 3.1X to 12.3X. (**C**) RAMP concentrator results in >95% recovery for all tested conditions (n=3 per concentration factor).

**Fig. 7.**
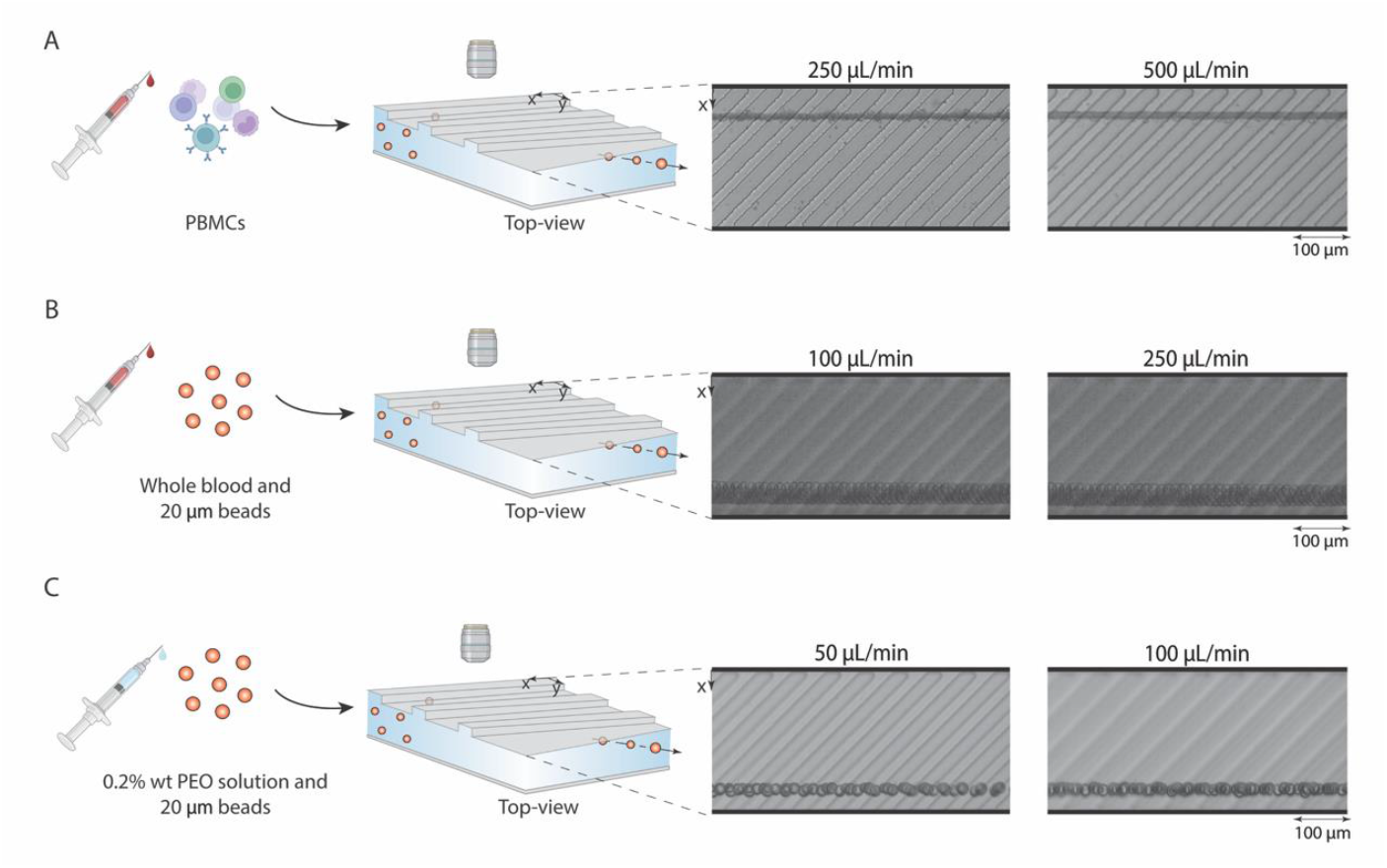
RAMP facilitates robust focusing across diverse sample types, including whole blood and viscoelastic fluids. (**A**) RAMP focuses peripheral blood mononuclear cells (PBMCs) in 0.2% F68 buffer. (**B**) RAMP can focus particles in undiluted whole blood. Here, we used 20 μm particles to visualize the focusing effect more clearly due to their bright fluorescence and clear visualization in high-speed imaging (**Movie S8**). (**C**) RAMP achieves viscoelastic focusing of particles in a polyethylene oxide (PEO) solution.

### Additional RAMP Applications

RAMP enables focusing across a range of sample types, including peripheral blood mononuclear cells (PBMCs) in buffer (**Figure 7A**), whole blood (**Figure 7B**), and viscoelastic polymer solutions (**Figure 7C**). Various liquid biopsy tests often use whole blood and PBMCs as the primary fluid (*22–24*). We show that the RAMP device can be used to align PBMCs at flow rates of 250 to 1000 μL/min (**Figure 7A** and **Supplementary Figure S8**), enabling the interrogation of clinically relevant human samples.

Another essential factor to consider in sample preparation is the ability to focus cells or particles in non-diluted whole blood. Due to the excessive number of RBCs in the blood (5 billion RBCs/mL), hydrodynamic particle-particle interactions disrupt equilibrium positions within the channel (*1*). To determine if particle-particle interactions impact RAMP’s focusing mechanism, we combined 20 μm particles with whole blood and flowed this suspension through the device at 100 and 250 μL/min flow rates. These bright, large fluorescent particles allowed us to directly visualize particle focusing in whole blood despite the high density of RBCs. The 20 μm particles in whole blood were still aligned near the channel sidewall, though at the opposite wall (near *x = W*) (**Figure 7B, Movie S8**) than the previously observed results with Newtonian fluids (**Figure 3E**). This flip in the focusing position is likely due to viscoelastic focusing resulting from the shear-thinning rheological properties of whole blood, discussed in the next section. The ability to directly focus particles or cells in undiluted whole blood is challenging for other inertial focusing methods and shows a remarkable property of the RAMP system, which may enable higher cellular throughput and direct sorting from whole blood. Previously reported methods dilute blood 10-fold to 25-fold before using inertial focusing (*24, 25*).

To test the role of viscoelastic forces in shifting the focusing position, we spiked the same 20 μm diameter particles in 0.2 wt% aqueous solution of polyethylene oxide (PEO), which imparts shear-thinning viscoelastic properties to the fluid (*27, 28*), and tested the solutions at 50 and 100 μL/min through the RAMP channel. Viscoelastic fluids have properties of viscous liquids and elastic solids, which cause them to flow but store energy when they are deformed, creating elastic stresses.

The combination of inertia and elasticity when using a viscoelastic fluid induces microparticle focusing by introducing an elastic lift force on particles (*26, 27*) (**Equation S5, Figure S9A**), arising from the imbalanced first normal stress difference across the channel, and it is highly dependent on particle size. In a rectangular channel, away from the top and bottom walls, elastic forces act towards the channel’s centerline for a shear-thinning fluid (**Figure S9B**). The elasticity effect is greater at higher PEO concentrations, flow rates, and particle confinement ratios (*26*).

Deborah’s number (*De*) is defined as the ratio of fluid relaxation time to characteristic time of the deformation process (*30, 31*) (**Equation S6**). A medium will exhibit fluid-like behavior when the characteristic time is large or the relaxation time is small, and solid-like behavior if the characteristic time is small or the relaxation time is large (*30*). Inertial effects can be quantified using the Reynolds number (**Equation S7**). Elastic and inertial effects can be compared through the Elasticity number (**Equation S8**), the ratio of elastic to inertial forces, where elasticity dominates when the Elasticity number is greater than 1. The Deborah number, Reynolds number, and the Elasticity number for each flow condition tested are detailed in **Table S6**. Estimates for viscosity and relaxation time were derived from published literature for the aqueous 0.2 wt% PEO solution (*29, 30*). Since the Elasticity number is greater by over two orders of magnitude at all flow rates (117.0), elastic forces dominate over inertial forces, confirming viscoelastic focusing as the primary focusing mechanism. In this regime, particles are concentrated toward the centerline of the rectangular channel (**Figure S9B**), where the transverse flow field is directed towards the far end of the ridge (**Figure S9C**), causing particles to be swept and focused at a position closer to the distant side wall (near *x = W*). At this position, elastic force (*F*_*E*_) balances the drag force (*F*_*D*_) and shear-gradient lift force (*F*_*SG*_) (**Figure S9D** and **Equation S9**). We limited testing to 100 μL/min as PDMS channels start to inflate at higher flow rates with highly viscous viscoelastic fluids due to increased pressure drop. Here, it is evident that particles retain the precise focusing effect of RAMP in viscoelastic fluids, and just as in whole blood, the particle focus position is observed at the far end of the ridge, proving that the focusing in undiluted whole blood is mainly controlled by viscoelastic forces (**Figure 7B**). Thus, RAMP is uniquely capable of achieving stable, deterministic focusing in undiluted whole blood samples by exploiting viscoelastic focusing effects.

## Discussion

Handling complex cellular biofluids involves aligning target cells into specific locations for isolation, characterization, or analysis to improve the sensitivity and efficiency of detection methods (*34*). For example, high-gradient magnetic sorting demands that cells be positioned near channel side walls, where magnetic gradients are strongest (*35*), and plasmonic or whispering-gallery mode lasers or sensors require cells to pass close to sensing features for effective signal generation (*36*). Similarly, imaging flow cytometry requires cells to be focused as tightly as possible in a single streamline in all three dimensions (*37*). Sample preparation of cells in biofluids is complicated by the cell size variability, which may vary across patients or within a single patient. Several biofluids, such as peritoneal and pleural fluids, also contain cellular debris and byproducts that can create channel blockages around pillars in the channel. Using rationally designed ridge arrangements, we have described a RAMP method that can focus particles and cells to any desired lateral position in a channel for a wide range of particle sizes, flow rates, and channel widths. We have leveraged these unique features to engineer a clog-free cell concentrator and to focus particles in undiluted whole blood.

Several microfluidic devices have been developed to accommodate and prepare diverse cell populations for downstream analyses (*1*). Active microfluidic manipulation techniques use external forces and fields, such as electric or magnetic fields, to focus particles (*34, 35*). Passive microfluidic manipulation techniques, including inertial focusing, can order particles more simply using channel microstructures. Microfluidic inertial focusing facilitates high-throughput particle control in a fluid stream using hydrodynamic inertial lift forces and secondary flow (*1*). Several microchannels have been designed for inertial focusing, such as serpentine, reverse wavy, contraction-expansion, spiral, and slanted groove configurations (*34, 36–39*) (**Table S1**). These inertial focusing devices have been especially successful for cell enrichment and separation (*40, 41*) (**Table S1**). Still, they align particles in variable locations across the channel cross-section, creating a size-based dispersion. Overall, the particle equilibrium position cannot be controlled with any of these methods, offering limited flexibility in optimally aligning particles for downstream analysis.

RAMP enables three-dimensional particle focusing within a single stream without relying on co-flow. Though microfluidic systems that utilize sheath flow have been used (*12, 46, 47*) for confining cellular streams with buffer co-flow, sheath streams induce significant shear stresses, reduce throughput, and add complexity to the overall design (*47*). In the past, ridge microstructures have been used in several configurations to create inertially focused streamlines of particles (*43–45*). Some of these techniques can focus diverse particle populations across various flow rates (*11, 52*). However, these systems cannot control the particle focus position in a channel (*15*). Remarkably, these systems also use high non-dimensional ridge heights (*α* = *h/*2*H*) ranging from 25.3% to 47.5% (*14, 43, 46*) which differ from RAMP’s low non-dimensional ridge height (*α* = *h/*2*H*) of 5%.

In contrast, RAMP is an adaptable, sheathless, high-throughput technique that allows tunable alignment of polydisperse cells and particles. We show precise tuning of the focus position, where small (5 μm) and large shifts (≥15 μm) in focusing position can be achieved by shifting the ridge microstructure within the channel. RAMP’s precise focusing capabilities extend to a wide range of particle sizes and flow rate regimes. We observed focusing of 10 to 30 μm diameter particles within a range of 2% of the width of the channel (5 µm) and 10 μm particles within a range of 4% of the height of the channel (2 μm) in the *z*-direction. Unlike various inertial techniques that focus particles in two equilibrium positions in the *z*-direction (*19, 20*), RAMP precisely focuses particles to one *z*-position. Manipulating cells’ 3D focus position is especially advantageous for imaging-based cytometry to minimize event dropouts, where typically only 50% of cells are focused (*49*).

As RAMP’s focusing behavior is largely unaffected by channel width, it is highly compatible with integration across several platforms with flexibility for scale-up. Further, RAMP can effectively focus PBMCs and particles in undiluted whole blood and viscoelastic fluids, demonstrating its adaptability to a wide range of biological fluids. Leveraging RAMP’s ability to focus polydisperse particles across a range of flow rates and channel widths, we developed a clog-free cell concentrator. Volume reduction is essential in biofluid processing, particularly for large volume samples such as 100 mL leukapheresis products or 1000 mL ascites samples. These samples often contain cell debris, clots, and NETs that clog devices and wrap around pillars used in previously demonstrated concentrator designs (*21*). In contrast, RAMP concentrators utilize ridge microstructures with a low non-dimensional ridge height (*α* ≅ 0.05) and a predominantly featureless channel design, effectively eliminating clogging. Using this approach, we achieved concentration of polydisperse cells and clusters with >95% recovery across multiple concentration factors.

In summary, RAMP uniquely facilitates precise control over the lateral focusing position of cells in a microchannel, independent of flow rate and cell size. This capability allows effective focusing in whole blood and facilitates clog-free cellular concentrators. The development of RAMP provides an easy-to-integrate and scalable focusing method that advances cell sensing and sorting applications from complex biofluids.

## Materials and Methods

### Experimental Design

This paper presents a new approach for tunable and precise inertial focusing across wide particle size and flow rate conditions. Ridge microstructure geometric properties were optimized to focus a particle size range capturing the cells in biofluid patient samples (10 to 30 μm) at flow rates enabling high-throughput processing (250 to 1000 μL/min). Channel dimensions and fluid conditions were kept constant unless specified when different flow rates or particle sizes were tested. Flow and particle size conditions tested were repeated to ensure reproducibility. The primary outcome was particle focusing position; secondary outcomes included the standard deviation of the focusing position.

### Microfabrication

Standard lithography techniques were used to fabricate polydimethylsiloxane (PDMS) devices at Massachusetts General Hospital. A cast for PDMS devices was produced by applying SU-8 50 at 2650 rpm and SU-8 5 at 3000 rpm for 30 seconds to a prebaked silicon wafer. The objective target thicknesses for the channel layer and microstructures were 50 μm and 5 μm, respectively. Channels were produced by irradiating the SU-8 layer with 365 nm UV radiation using a Mylar mask as a stencil. Ridge microstructures were formed by treating the exposed SU-8 layer with Baker BTS-220 SU-8 developer. PDMS made using the Sylgard PDMS kit mixed at a 10:1 base-to-cross-linker ratio was poured into the SU-8 mold, degassed in a vacuum chamber, and cured in a convection oven at 65 °C for 12 hours. A 1.2 mm biopsy punch was used to drill inlets and outlets, and PDMS devices were plasma bonded to 3-inch by 1.5-inch glass slides and baked at 150 °C for 3 hours. This fabrication process creates a viewing window beneath the channel, so particle focusing in the *x*-*y* plane can be observed with an inverted microscope.

### Side-View Device Fabrication

To observe particle focusing in the *y*-*z* plane, a channel with ridge microstructures was bonded to a PDMS base layer. A viewing window is created by cutting close to the microchannel and base layer side wall, leaving a thin layer of PDMS enclosing the channel. To restore transparency in the viewing window, a thin layer of uncured PDMS was poured onto a glass slide, and the cut channel was adhered to it and baked at 80°C for 30 minutes. This fabrication process is illustrated in **Figure S10**.

### Particle and Cell Inertial Focusing in Microfluidic Flow

A Harvard Apparatus syringe pump and Tygon thermoplastic tubing were used to flow Fluoro-Max 10 μm, 15 μm, 20 μm, and Duke Standards 30 μm fluorescent particles at flow rates ranging from 250 to 1000 μL/min through devices. A Nikon Ti-U inverted microscope and Phantom 4.2, Vision Research Inc., and QImaging Retiga 2000R CCD cameras were used to capture high-speed and fluorescent videos of the particles and cells flowing in microchannels. COMSOL Multiphysics was used to simulate velocity fields.

### CTC Culture

The protocol used for MGH-BRx-142 CTC culture is reported in (*50*). MGH-BRx-142 CTCs were cultured in media with RPMI-1640, 1X B27 (Gibco), EGF (20 ng/mL) (Gibco), bFGF (20 ng/mL) (Gibco), and 1X antibiotic/antimycotic (Life Technologies) in ultralow attachment flasks (Corning). Then, several rounds of manual pipetting were done to dissociate cells. CTC culture was performed at 37°C, under hypoxic conditions (4% O_2_), and with 5% CO_2_.

### PBMC Collection

For experiments involving whole blood, blood was obtained from healthy internal donors or whole blood healthy donor samples from Research Blood Components, LLC. Experimental protocols were reviewed and authorized by the Massachusetts General Hospital to obtain informed consent for internal whole-blood donations under Protocol 2009-P-000295. The whole blood sample was aliquoted into 10 to 20 mL volumes in Vacutainer EDTA Tubes (BD Biosciences). Whole blood was diluted at a 1X working concentration with 0.2% F68 Buffer and processed through the CTC iChip operating at a pressure of 48.0 psi. A 0.2% F68 Buffer was used as co-flow through the CTC iChip. The non-equilibrium inertial separation array devices in the CTC-iChip will separate PBMCs from red blood cells and platelets into a product streamline containing clean 0.2% F68 Buffer. The resulting buffy coat is fixed for 10 minutes using 0.5% formaldehyde. The PBMC concentration was determined by assessing 100 μL of the product through an XP-300TM Automated Hematology Analyzer. More robust cell counts were subsequently done with disposable hemocytometers (Bulldog-Bio) and Nageotte hemocytometers (Hausser Scientific).

### Particle Viscoelastic Focusing in Microfluidic Flow

A 2000 ppm PEO solution was combined with Fluoro-Max 20 μm diameter particles to test RAMP’s viscoelastic focusing capabilities. High-speed and fluorescent microscopy videos of particles flowing in PEO were captured using the same methodology to observe particle inertial focusing.

### Image Analysis

Particle center of mass tracking was conducted with ImageJ software on all high-speed and fluorescent microscopy videos. The workflow follows this sequence: 1) cropping and adjusting the stack of frames to show just outside the channel boundary, 2) reduction of the stack (by a factor of ∼3) to ensure particles are counted once, 3) inversion of the stack, 4) subtract an average intensity frame of the stack from each frame of the main video, 5) adjust brightness and contrast to enhance particle visibility while minimizing noise, 6) convert the video to 8-bit and binary form, 7) adjust the threshold for optimal contrast between particles and background, 8) apply erosion and dilation to the particles to refine outlines and eliminate residual noise, and 8) determine particle center of mass positions using ImageJ’s *Analyze Particles* feature. Following particle analysis on ImageJ, the resulting file with the particle area and the center of mass in the *x*- and *y*-directions was processed for statistical analysis. Particles with areas smaller than 20 pixels are artifacts from the erosion and dilation steps and were omitted from analysis.

### Statistical Analysis

Particle positional data were analyzed with Python Pandas and Numpy packages. For each group corresponding to a flow rate condition or particle size tested, outliers were removed using the interquartile range (IQR) method using a threshold of 1.5xIQR to ensure that image artifacts are not included in the analysis. The mean focusing position and its standard deviation were calculated.

## Funding

National Cancer Institute grant U01CA214297 (MT)

National Cancer Institute grant U01CA268933 (MT)

National Cancer Institute grant R01CA255602 (MT, AM)

National Heart, Lung, and Blood Institute grant K25HL169816 (AM)

National Science Foundation Graduate Research Fellowship award 2141064 (APM)

Disclaimer: Opinions, findings, conclusions, or recommendations in this manuscript are those of the author and do not necessarily reflect the views of the National Science Foundation.

## Author contributions

Conceptualization: APM, AM, MT

Methodology: APM, AM, SRB, VK

Investigation: APM, VRP

Visualization: APM, AM

Supervision: APM, AM, MT

Writing—original draft: APM, AM, SRB

Writing—review & editing: APM, AM, SRB, VRP, QC, VK

## Competing interests

All authors’ interests were reviewed and managed by Massachusetts General Hospital and Mass General Brigham in accordance with their competing interests policies.

## Data and materials availability

All data needed to evaluate the conclusions in the paper are present in the paper and/or the Supplementary Materials. Additional data related to this paper may be requested from the authors.

## Notes

### Competing Interest Statement

The authors have declared no competing interest.

### Summary of Updates

author contributions updated for Varun Kulkarni

