## Supplementary File for "Ridge-assisted Micro Positioning of Cells and Particles in a Microchannel"

Adriana Payan-Medina *et al.*

**This PDF file includes:**

Supplementary Text
Figs. S1 to S8
Tables S1 to S6
Movies S1 to S8
References (1 to 13)

**Other Supplementary Materials for this manuscript include the following:**

Movies S1 to S8

### Supplementary Text

<u>Chevron Theory for Inertial Focusing</u>

Trapping in ridge-assisted microfluidic devices is governed by a balance of secondary-flow-induced drag and lift forces. Since trapping is occurring away from the top and side walls, we can approximate the lift force as $F_L = C_L \rho U^2 a^4 / H^2$, where $C_L$ is the lift coefficient, $\rho$ is the fluid density, $U$ is the average velocity, $a$ is the particle diameter, and $H$ is the height of the channel. Theoretical treatment of secondary flow induced by ridges has been presented in (*1*). Following their analysis for thin channels ($W >> H$) and shallow grooves ($h/H << 1$), the drag force on a particle can be written as $F_D = 3\pi\mu a U_{y1}$. Thus, the ratio of lift to drag force can be written as **Equation S1**.

**Equation S1.**

$$\frac{F_L}{F_D} = \frac{\frac{C_L \rho U^2 a^4}{H^2}}{3\pi\mu a U_{y1}}$$

**Equation S2.**

$$U_{y1} = 6\alpha^2 U \left(\frac{3}{2}\frac{(H-z)(z)}{H^2} - \frac{z}{2H}\right)\left(\frac{2\pi H}{\lambda} - 1\right) \mathrm{Cos}\,\theta$$

In **Equation S2,** λ is the wavelength, $\theta$ is the angle of the ridges, and $\alpha$ is a geometrical nondimensional number. A relation for ridge height ($h$) is presented as $h = 2\alpha H$. Thus, the lift-to-drag force ratio can be approximated as given in **Equation S3**.

**Equation S3.**

$$\frac{F_L}{F_D} = \frac{C_L \rho U}{18\pi\mu\alpha^2 \left(\frac{3}{2}\frac{(H-z)(z)}{H^2} - \frac{z}{2H}\right)\left(\frac{2\pi H}{\lambda} - 1\right) \mathrm{Cos}\,\theta} a^3/H^2$$

Assuming that trapping is often seen at $z \cong \frac{3H}{4}$, this equation can be reduced to a more simplified form (**Equation S4**).

**Equation S4.**

$$\frac{F_L}{F_D} \cong \frac{8}{27\pi^2}\frac{C_L \rho U \lambda}{\mu\alpha^2 \mathrm{Cos}\,\theta}\left(\frac{a}{H}\right)^3$$

Though these results are approximate solutions derived through various assumptions, and the slip velocity will be higher than the vertical component of velocity, these results show that among a few parameters, particle trapping will be strongly affected by confinement ($a/H$) and $\alpha$. Based on this analysis, we designed devices with low $\alpha$ and high confinement ratios ($a/H > 0.2$). In this analysis, $C_L = 0.12$, based on the work done in (*2*).

Elastic lift force: Elastic lift force is proportional to normal stress ($N_1$) variation over the particle volume ($a^3$). $C_e$ represents the elastic lift coefficient.

**Equation S5.**

$$F_e = C_e a^3 \nabla N_1$$

Deborah's number ($De$): $De$ is a ratio of the characteristic time of the fluid ($\tau$) and the characteristic time of the deformation process ($t_f$).

**Equation S6.**

$$De = \frac{\tau}{t_f} = \frac{\tau Q}{H^3}$$

Reynolds number ($Re$): $Re$ is used to compare fluid inertial and viscous forces. Fluid density, average velocity, channel hydraulic diameter, and viscosity are $\rho$, $U$, $D_h$, and $\mu$, respectively.

**Equation S7.**

$$Re = \frac{\rho U D_h}{\mu}$$

Elasticity number ($El$): $El$ is a ratio of Deborah's number divided by Reynolds number and can be used to compare a fluid's elastic and inertial properties.

**Equation S8.**

$$El = \frac{De}{Re}$$

The ratio of elastic lift force and drag force can be written as Equation S9, denoting a weaker dependence on particle size ($\propto a^2$) than the ratio of inertial lift force and drag force ($\propto a^3$).

**Equation S9.**

$$\frac{F_e}{F_D} = \frac{C_e a^3 \nabla N_1}{3\pi\mu a U_{y1}} = \frac{C_e a^2 \nabla N_1}{3\pi\mu U_{y1}}$$

Computation Modelling

Steady state 3D fluid flow simulations were conducted in COMSOL Multiphysics using the Navier–Stokes equation. A schematic diagram of the boundary conditions is shown in Figure S1, below. A velocity inlet boundary condition was applied at the inlet, an atmospheric pressure outlet boundary condition was applied, and a no-slip boundary condition was imposed on the rest of the surfaces. A free tetrahedral mesh with a minimum element size of 0.5 µm was used. We conducted a mesh refinement study to ensure that the results were independent of mesh element size.

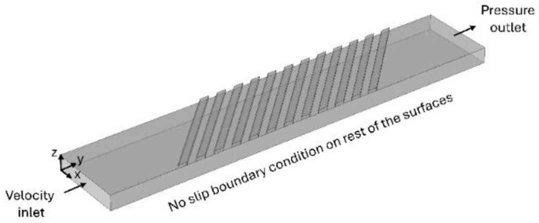


**Figure S1.** Boundary conditions used in the fluid flow simulation.

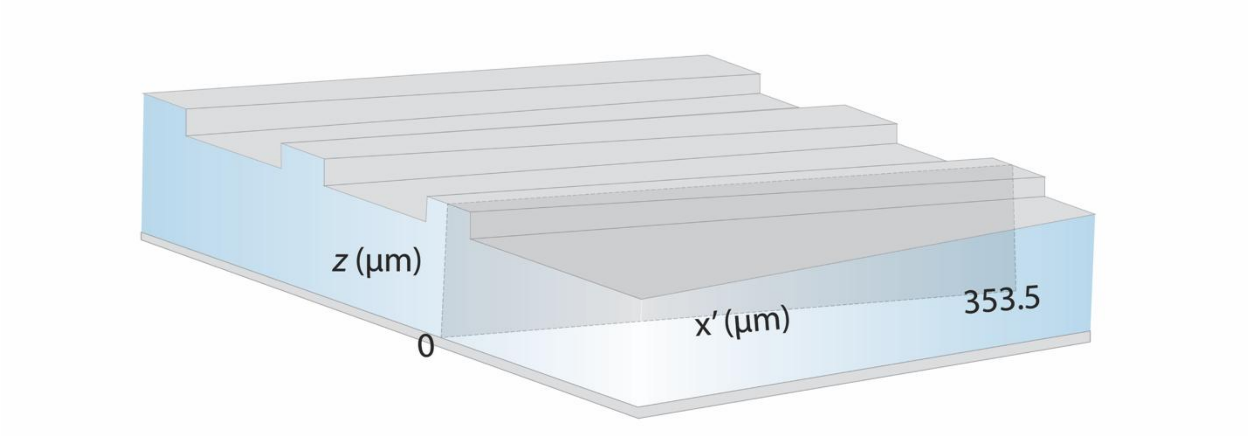


**Figure S2.** A schematic representation of a RAMP device shows the cross-section parallel to the ridge microstructures used for presenting the velocity field and the microvortex.

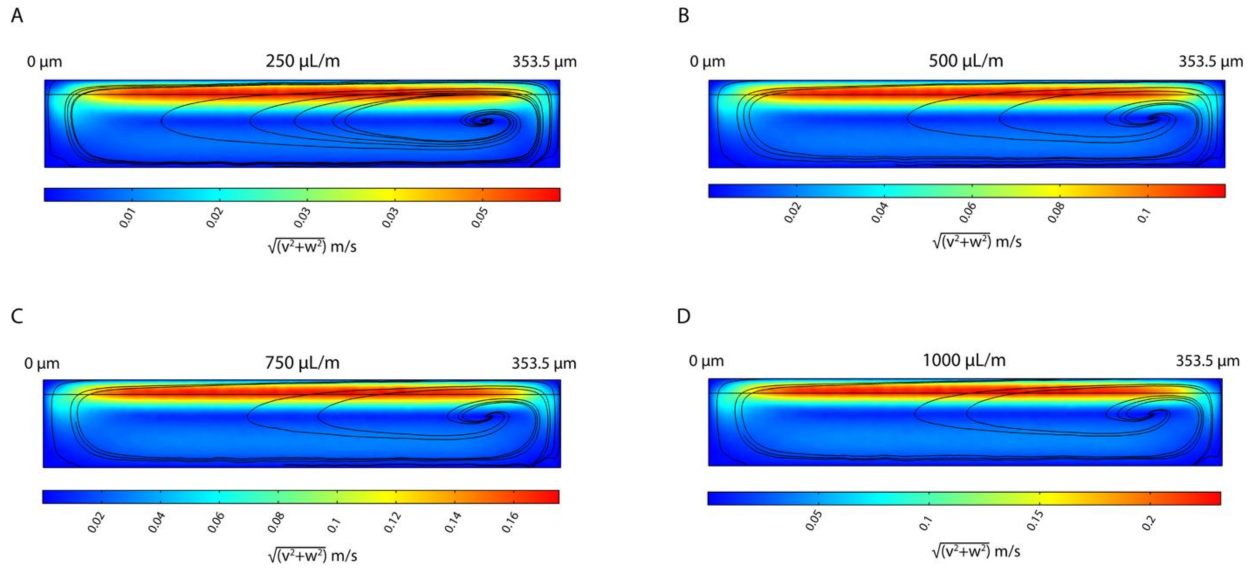


**Figure S3.** Finite element simulations of fluid flow in a RAMP device show equivalent recirculation regions formed at several flow rates. Simulations were performed at flow rates of (**A**) 250 µL/min, (**B**) 500 µL/min, (**C**) 750 µL/min, and (**D**) 1000 µL/min.

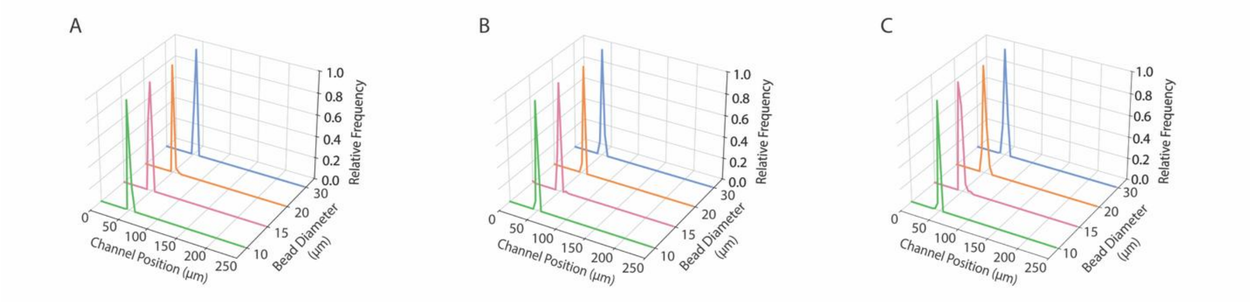


**Figure S4. RAMP aligns 10, 15, 20, and 30 μm particles across a wide range of flow rates.** Histograms of particle centroid positions are shown for flow rates of (**A**) 500 μL/min, *Re* = 50.4 (n=118 (10 μm), 99 (15 μm), 146 (20 μm), 105 (30 μm) particles), (**B**) 750 μL/min, *Re* = 75.6 (n=180 (10 μm), 99 (15 μm), 71 (20 μm), 143 (30 μm) particles), and (**C**) 1000 μL/min, *Re* = 100.8 (n=132 (10 μm), 99 (15 μm), 165 (20 μm), 163 (30 μm) particles).

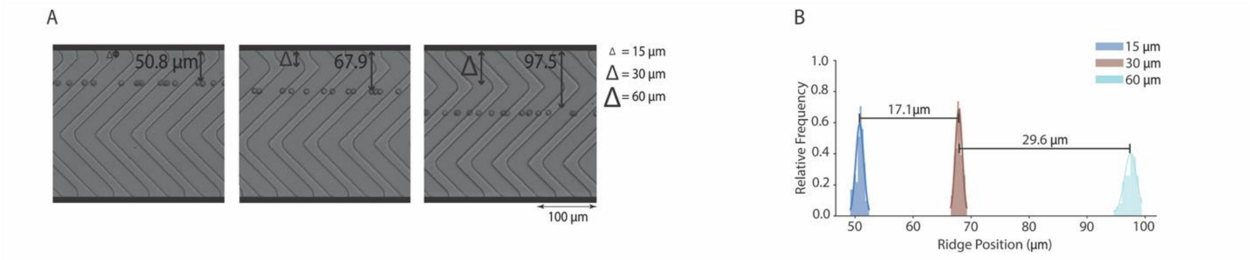


**Figure S5. Large focusing position shifts in 10 µm particle streamlines are achieved with proportional shifts in the position of center ridges.** (**A**) Shifts in particle focus position are observed with high-speed camera streak images as the central ridge structure is shifted in steps of 15 µm and 30 µm. (**B**) A frequency distribution plot of particle centroid positions shows the lateral streamline shift that occurs as the center ridge is shifted (15 µm (n = 131), 30 µm (n = 83), and 60 µm (n = 105)).

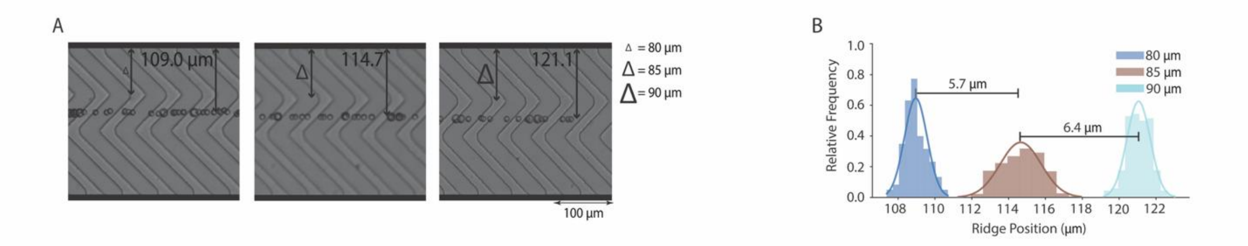


**Figure S6. Focusing position shifts in a mixture of 10 μm and 15 μm particles are achieved with proportional shifts in the position of center ridges.** (**A**) Shifts in particle focus position are observed with high-speed camera streak images as the position of the central ridge structure is shifted in steps of 5 μm from the channel side wall. (**B**) A frequency distribution plot of particle centroid positions describes the lateral streamline shift that occurs as the center ridge is shifted 80 μm (n = 384), 85 μm (n = 403), and 90 μm (n = 191) from the channel sidewall.

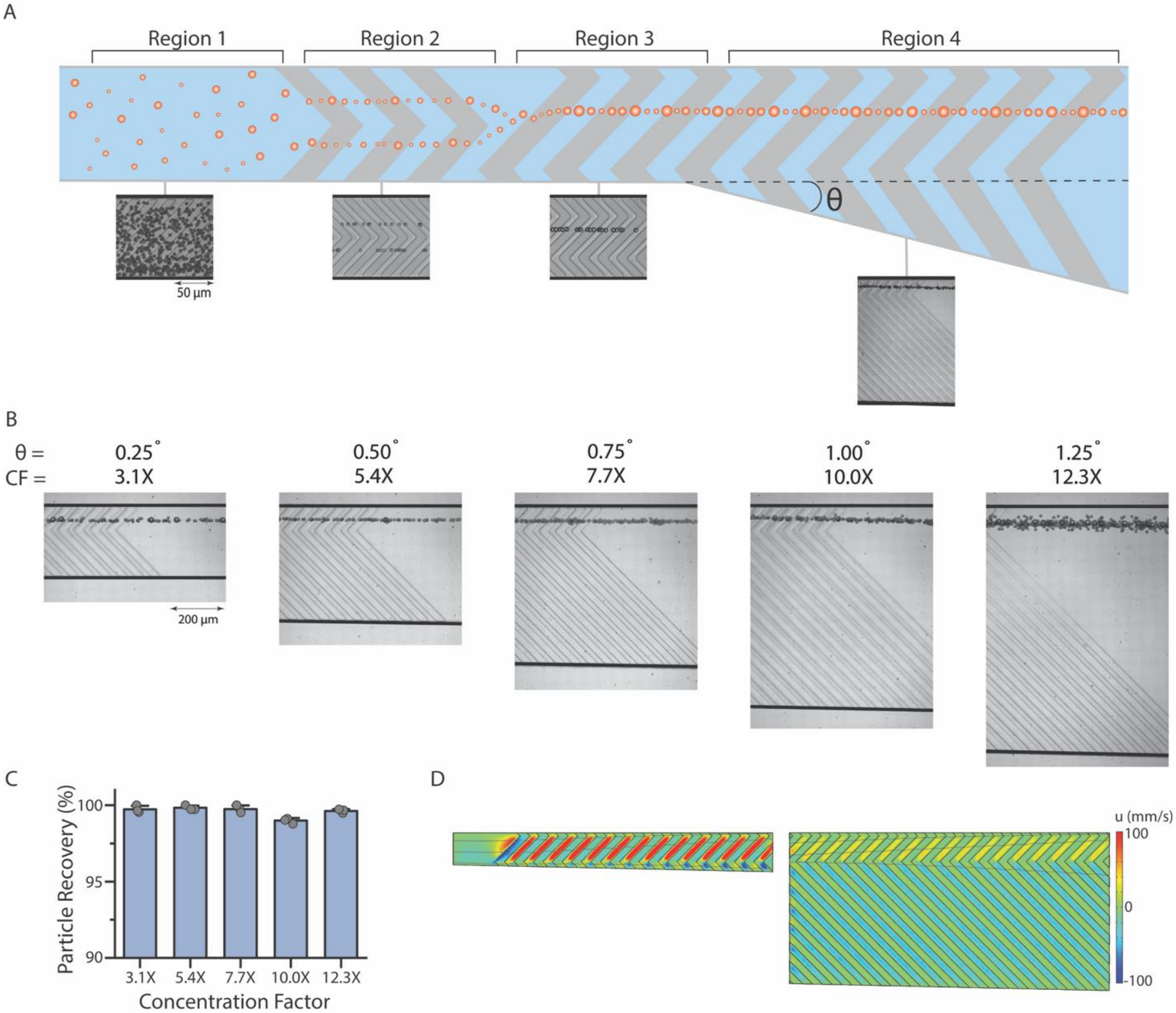


**Figure S7. A RAMP device is used to successively align and concentrate polydisperse particles.** (**A**) High-speed streak imaging showing the organization of polydisperse particles in various regions. (**B**) High-speed microscopy imaging shows a polydisperse mixture of particles (10 and 20 μm diameters) being concentrated with concentration factors (CFs) of 3.1X, 5.4X, 7.7X, 10.0X, and 12.3X. (**C**) RAMP cell concentration results in recovery greater than 99% for each CF tested (n = 3 replicates per concentration factor). High-speed videos show the polydisperse particle mixture flowing through concentration factors of 3.1X and 12.3X (**Movie S4** and **Movie S5**). (**D**) Flow simulation results show a change in the transverse velocity profile as fluid siphoning is implemented. Gray lines show streamlines.

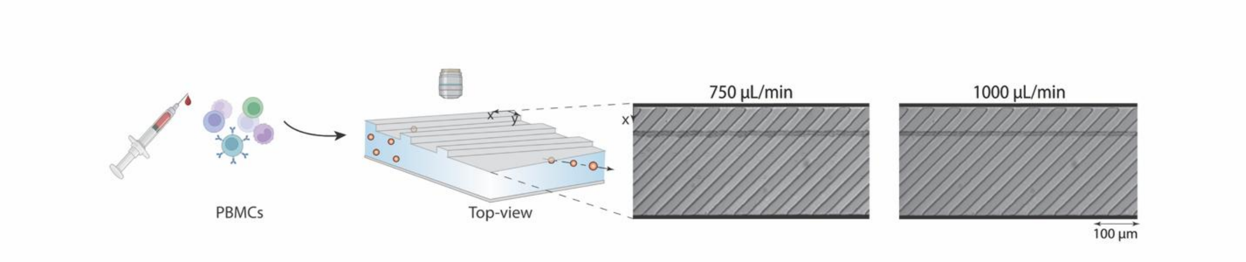


**Figure S8.** RAMP focuses peripheral blood mononuclear cells (PBMCs), showing its utility for clinically relevant cell suspensions.

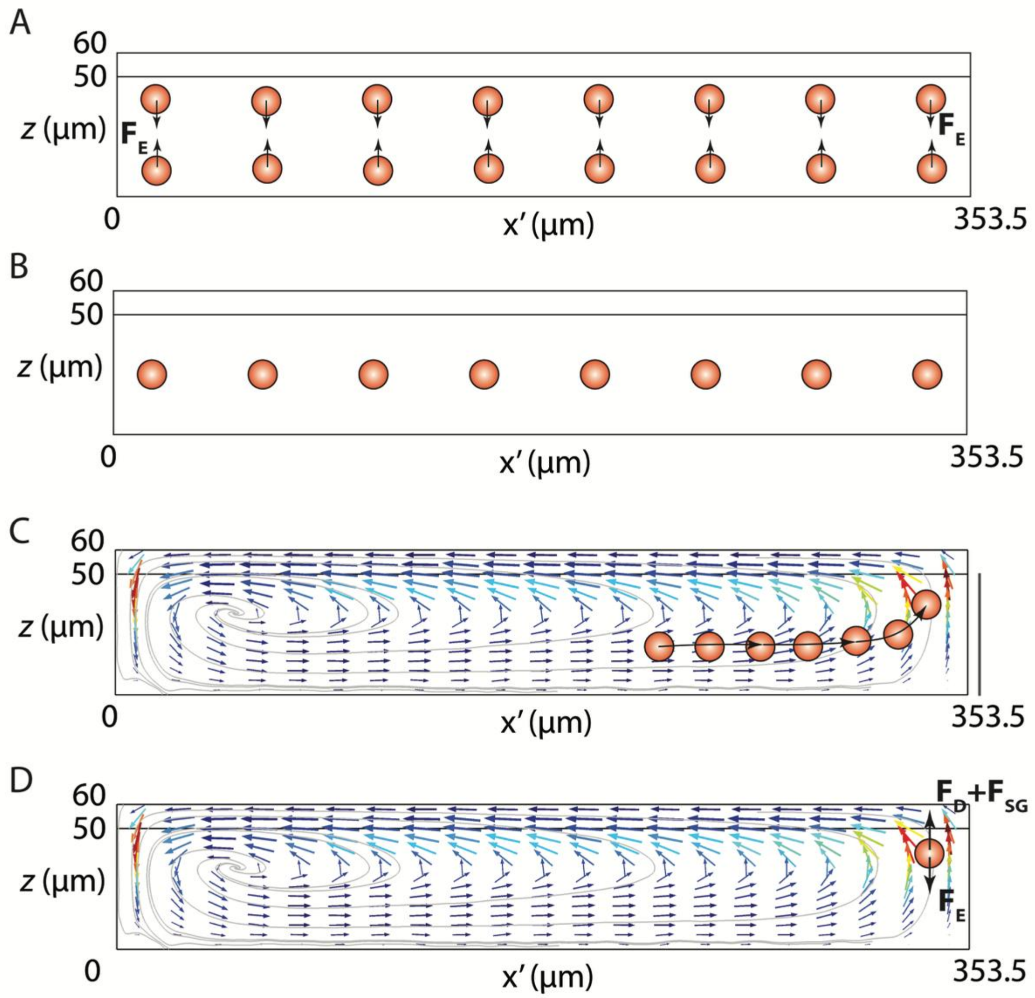


**Figure S9. Schematics of viscoelastic focusing.** (**A-B**) For the conditions tested in this work, elastic forces ($F_E$) dominate inertial forces; as a result, particles experience an elastic force towards the center of the rectangular channel. (**C-D**) Secondary flow produced by ridges sweeps particles towards the distant side wall of the channel, where elastic force towards the center of the channel balances the drag force due to vortex ($F_D$) and shear-gradient inertial lift force ($F_{SG}$).

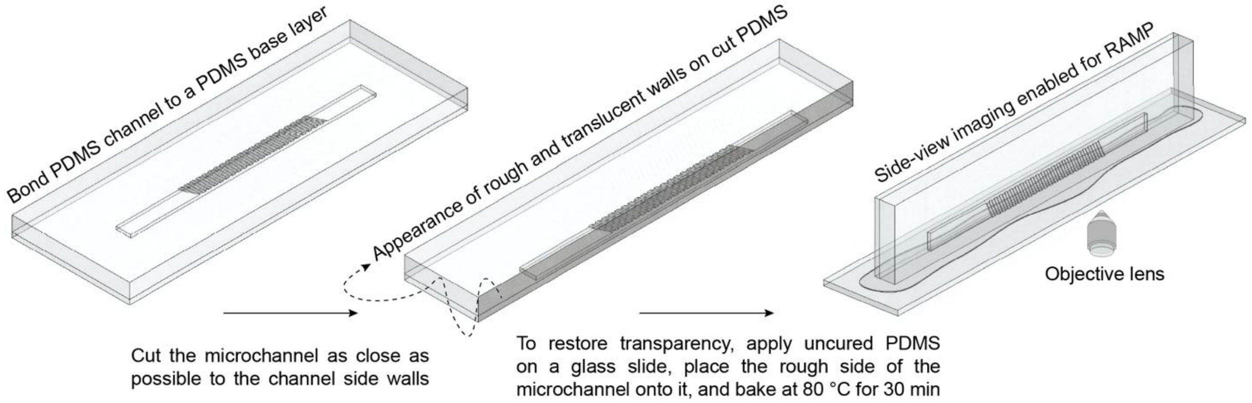


**Figure S10. Fabrication process for PDMS device with a side-view (*y*-*z* plane) window.**

**Table S1.** An overview of microchannel geometries used for inertial focusing with corresponding flow conditions, applicable particle size ranges, and focusing behavior.

| **Channel Type** | **Channel Visual** | **Tunable focusing** | **Size independent focusing** | **Flow Rate independent focusing** | **Focusing in height (Z direction)** |
|---|---|---|---|---|---|
| Current Study (RAMP) | 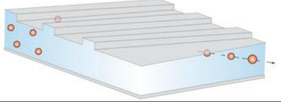 | Yes | Yes | Yes | Single line |
| Serpentine and curvilinear | 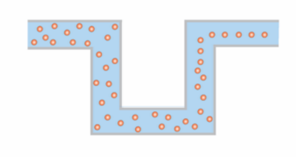 | No | No | No | Double line |
| Obstacled curvilinear (*3–7*) | 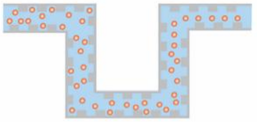 | No | No | No | Double line |
| Asymmetric serpentine (*8*) | 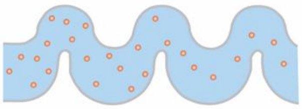 | No | No | No | Double line |
| Spiral (*9–12*) | 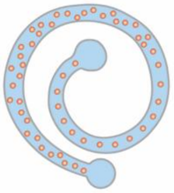 | No | No | No | Double line |
| Obstacled-spiral (*13–15*) | 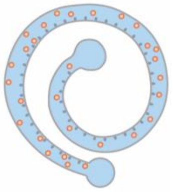 | No | Yes | Yes | Double line |
| Contraction-expansion (*16–18*) | 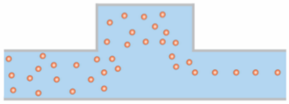 | No | No | No | Double line |
| Reverse wavy (*19*) | 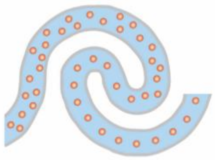 | No | No | No | Double |
| Straight microchannel with rectangular cross-section (*20*, *21*) | 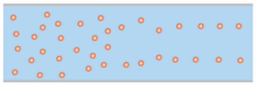 | No | No | No | Double or multiple |

| | | | | | |
|---|---|---|---|---|---|
| Straight microchannel with triangular cross-section (*22–24*) | 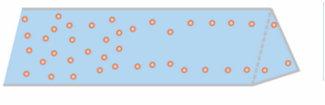 | No | Yes (in some cases) | Yes (in some cases) | Single or double |
| Mixed channel cross-section in a straight channel (*25*) | 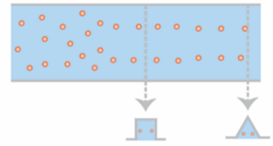 | Yes (to some extent) | No | No | Single |
| Stepped straight microchannel (*26*) | 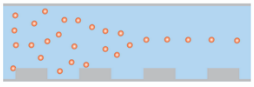 | No | Yes | Yes | Single |
| Grooved microchannels (*27–29*) | 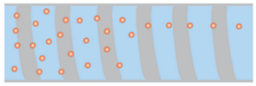 | No | Yes | Yes | Single |

**Table S2.** 10, 15, 20, and 30 µm particles' focus positions and standard deviations at flow rates of 250 to 1000 µL/min captured using high-speed microscopy imaging.

| **Particle Size (µm)** | **Focus Position (µm)** | | | |
|---|---|---|---|---|
| | **250 µL/min Re = 25.2** | **500 µL/min Re = 50.4** | **750 µL/min Re = 75.6** | **1000 µL/min Re = 100.8** |
| 10 | 57.8 ± 0.7 | 55.0 ± 0.6 | 52.2 ± 0.5 | 54.0 ± 0.5 |
| 15 | 54.2 ± 1.3 | 51.1 ± 0.9 | 51.3 ± 5.2 | 51.3 ± 2.8 |
| 20 | 54.4 ± 0.9 | 54.4 ± 1.3 | 56.2 ± 0.9 | 54.0 ± 3.2 |
| 30 | 59.5 ± 2.3 | 56.5 ± 1.8 | 53.8 ± 1.8 | 53.1 ± 2.6 |

**Table S3.** Focusing positions of 10 μm particle in *z*-direction at flow rates ranging from 250 – 1000 μL/min captured using fluorescence microscopy in *y*-*z* plane (side-view).

| Flow Rate (μL/min) | Mean Peak Position (μm) |
|---|---|
| 250 | 30.6 ± 0.6 |
| 500 | 31.5 ± 0.6 |
| 750 | 31.5 ± 0.6 |
| 1000 | 32.4 ± 0.5 |

**Table S4.** Focusing positions of 10 μm particles for small and large focus position shifts at a flow rate of 500 μL/min and *Re* of 50.4 captured using a high-speed camera.

| **Ridge Position (μm)** | **Focus Position (μm)** | |
|---|---|---|
| | **10 μm Particles** | **10 and 15 μm Particles** |
| 15 | 50.8 ± 0.7 | |
| 30 | 67.9 ± 0.6 | |
| 60 | 97.5 ± 1.0 | |
| 80 | 109.1 ± 0.6 | 109.0 ± 0.6 |
| 85 | 113.9 ± 0.5 | 114.7 ± 1.1 |
| 90 | 119.8 ± 0.6 | 121.1 ± 0.6 |
| 95 | 126.7 ± 0.6 | |

**Table S5.** Average cell and particle recovery in RAMP concentrators (n=3).

| Concentration Factor | Cell Recovery (%) | Particle Recovery (%) |
|---|---|---|
| 3.1X | 97.6 ± 1.1 | 99.7 ± 0.2 |
| 5.4X | 99.5 ± 0.2 | 99.8 ± 0.1 |
| 7.7X | 98.4 ± 0.2 | 99.7 ± 0.2 |
| 10.0X | 95.1 ± 0.5 | 99.0 ± 0.1 |
| 12.3X | 96.8 ± 0.8 | 99.6 ± 0.1 |

**Table S6.** Deborah's number (*De*), Reynold's number (*Re*), and the Elasticity number (*El*) calculations for flow rates of 50 and 100 μL/min.

| Flow Rate (μL/min) | *De* | *Re* | *El* |
|---|---|---|---|
| 50 | 130.0 | 1.111 | 117.0 |
| 100 | 260.0 | 2.222 | 117.0 |

**Movie S1.**
Top-view of 10 µm particles flowing at 500 µL/min through a channel with inclined straight ridges.

**Movie S2.**
Top-view of 10 (left) and 30 (right) µm particles flowing at 500 µL/min through a channel with inclined straight ridges.

**Movie S3.**
Top-view of 10 µm particles flowing at 500 µL/min through a channel with center ridge structures at 80 µm from the channel side wall (left) and 90 µm from the channel side wall (right).

**Movie S4.**
Top-view of 9, 10, and 20 µm particles flowing through a RAMP concentrator with a concentration factor of 3.1X and operating pressure of 20 psi.

**Movie S5.**
Top-view of 9, 10, and 20 µm particles flowing through a RAMP concentrator with a concentration factor of 12.3X and operating pressure of 20 psi.

**Movie S6.**
Top-view of white blood and circulating tumor cells flowing through a RAMP concentrator with a concentration factor of 3.1X and operating pressure of 15 psi.

**Movie S7.**
Top-view of white blood and circulating tumor cells flowing through a RAMP concentrator with a concentration factor of 12.1X and operating pressure of 15 psi.

**Movie S8.**
Top-view of 20 µm particles flowing at 250 µL/min in whole blood through a device with RAMP.
